# Rotten-skin disease significantly changed giant spiny frog(*Paa spinosa*) gut microbiota

**DOI:** 10.1101/2020.01.13.905588

**Authors:** Tuoyu He, Yun Jiang, Pengpeng Wang, Jianguo Xiang, Wangcheng Pan

**Author notes:** Correspondence author: jianguo Xiang, College of Animal Science and Technology, Hunan Agricultural University, Changsha, 410125,China.

## Abstract

The composition and abundance of gut microbiota is essential for host health and immunity. Gut microbiota is symbiotic with the host, so changes in the host diet, development, and health will lead to changes in the gut microbiota. Conversely, changes in the gut microbiota also affect the host conditions. In this experiment, 16S rRNA high-throughput sequencing was used to compare the gut microbiota composition of 5 healthy *Paa Spinosa* and 6 *P. spinosa* with rotten-skin disease. Results: the gut microbiota composition was significant difference between diseased *P. spinosa* and the healthy *P. spinosa*; LEfSe analysis showed that the relative abundance of *Methanocorpusculum, Parabacteroides, AF12, PW3, Epulopiscium*, and *Oscillospira* were significantly higher in the diseased *P. spinosa*, while the relative abundance of *Serratia, Eubacteium, Citrobacter*, and *Morganella* were significantly lower. Conclusion: Rotten-skin disease changed *P. spinosa* gut microbiota significantly; The relative abundance of *Epulopiscium* and *Oscillospira* might be related to the health conditions of the host skin and gallbladder; The relative abundance of *Serratia* and *Eubacteium* might be important for maintaining the gut microbiota ecosystem.

## Introduction

The giant spiny frog (*Paa spinosa*) is a large edible frog distributed in the mountains of China and Vietnam (1). It is favored by people for its great economic and medicinal value (2). In recent years, due to the increase of market demand and the destruction of habitats, the wild *P. spinosa* have been declined sharply. The Chinese Red Animal List has listed *P. spinosa* as a “vulnerable” species(3) (4). The artificial breeding has provided an effective way to meet market demand and protect wild *P. spinosa*. However, frequent disease problems seriously restrict the development of the *P. spinosa* industry (1) (5).

The rotten-skin disease is a common disease of *P. spinosa*, which is characterized by a dull epidermis and white spots appearing at the beginning, and then the epidermis falls off and begins to rot until the bone is exposed(6). Many pathogens cause rotten-skin disease, such as *Proteus mirabilis* and *Yersinia kristensenii*(6). Some diseased frogs will not die immediately, but growth is affected (7), some pathogens not only cause diseased frogs to show signs of rot but also cause a large number of deaths, such as *Chytridiomycosis*. The diversity of pathogens of rotten-skin disease bring difficulties to routine methods of preventing the disease, a new method is needed immediately.

Animal gut microbial communities as host “microbial organs” play an important role in host immunity and health, such as promoting the absorption of nutrients(8) (9) (10), impeding pathogens colonization in the gut (11) and regulating host immunity to maintain host health. Gut microbiota is symbiotic with the host, so changes in the host diet, development, and health can lead to changes in the gut microbiota (12)(13)(14). Conversely, changes in the gut microbiota also affect the host conditions. The normal microbiota composition is the guarantee for maintaining the physiological function of the host (15). When the physiological function of the host is abnormal (such as the disease involving), the gut microbiota composition will change as well(16). In recent years, a new understanding of aquatic animal diseases has been gained by comparing the gut microbiota of diseased aquatic animals with healthy ones (Table 1). So we hypothesized that gut microbiota changed significantly between the healthy *P. spinosa* and the rotten-skin diseased *P. spinosa*. And comparing the composition of gut microbiota of healthy *P. spinosa* and the rotten-skin diseased *P. spinosa* we will know the microbiota change, which may be vital to discover the antagonistic bacteria of *P. spinosa* rotten-skin disease and explore new methods to prevent and control rotten-skin disease.

**Table 1.** Recent studies about gut microbiota between healthy and diseased samples

| Species | Disease | Finding | Authors |
| --- | --- | --- | --- |
| Grass carp<br>(*Ctenopharyngodon idellus*) | Enteritis | The association between changes of the gut microbiota and enteritis in grass carp | (39) |
| Crucian Carp<br>(*Carassius auratus*) | Red-Operculum Disease | The surge of some potential pathogens as bacterial signatures that were associated with “red-operculum” disease in crucian carps | (16) |
| largemouth bronze gudgeon<br>(*Coreius guichenoti*) | Furunculosis | The presence of healthy carriers of pathogenic *Aeromonas salmonicida* among the farmed fish, and the gut appeared as a probable infection source for furunculosis in largemouth bronze gudgeon. | (37) |
| Ayu<br>(*Plecoglossus altivelis*) | *Vibrio anguillarum* infection | *Vibrio anguillarum* infection substantially disrupted the compositions and interspecies interaction of ayu gut bacterial community. | (38) |
| Zebrafish(*Barc hydanio rerio* var) | *Aeromonas hydrophila* infected | the invasion of pathogen could change the gut microbiota composition and induce gut innate immune responses in zebrafish | (51) |
| Gibel Carp<br>(*Carassius gibelio*) | *Cyprinid herpesvirus 2* (CyHV-2) Infection | The composition was dramatically altered following CyHV-2 infection ; *Plesiomonas* was highly abundant in infected samples, and could be used as a microbial biomarker for CyHV-2 infection | (52) |
| Chinese mitten crab (*Eriocheir sinensis*) | White spot syndrome virus (WSSV) infection | Changes in gut microbiome were closely associated with the severity of WSSV infection and that indicator taxa could be used to evaluate the crab health status. | (53) |
| Shrimp<br>(*Litopenaeus vannamei*) | Acute hepatopancreatic necrosis disease | Shrimp health is highly relevant to the homeostasis of its gut bacterial community. | (35) |

In this paper, the 16S rRNA amplicon high-throughput sequencing technology was used to investigate the effects of rotten-skin disease on gut microbiota composition in *P. spinosa*. The potential probiotics and antagonistic bacteria were screened out in the healthy *P. spinosa* through comparing the composition of gut microbiota between healthy and diseased *P. spinosa*, and expected to enrich the theories about regulating gut microbiota structure by using microbial ways and realizing the microecological prevention of rotten-skin disease in *P. spinosa*.

## Materials and methods

### Sample Collection

All animal experiments were conducted in accordance with the recommendations in the Guide for the Care and Use of Laboratory Animals of the National Institute of Health(NIH). The experimental animals were approved by the experimental animal ethic committee of Hunan Agriculture University. All samples were collected from Shimen County, Hunan ProvinceWeixin Town *P*.*spinosa* farm (110°29’-111°33’E, 29°16’-30°08 N). There were 11 *P*.*spinosa* in the experiment, including 5 Healthy *P*.*spinosa* (H) and 6 diseased *P*.*spinosa* (D). The frog was anesthetized and dissected under sterile conditions, collected gut contents, transferred to 2 ml sterile EP tubes and stored at −80 ° C, used for subsequent DNA extraction.

### DNA extraction and high-throughput sequencing

The gut microbiota DNA was extracted using a DNeasy PowerSoil Kit (QIAGEN, Germany). The V4-V5 hypervariable region of the prokaryotic 16S rRNA gene was amplified using the universal primer pair 515F and 909R, with a 12-nt sample-specific barcode sequence, including at the 5’-end of the 515F to distinguish samples (17) (18). PCR was performed, and amplicons were sequenced using a MiSeq system at Guangdong Meilikang Bio-Science, Ltd. (China), as described previously (1). The raw sequences were merged using FLASH-1.2.8 software and processed using the QIIME pipeline 1.9.0 as described previously(1)(19). Chimeric sequences were identified and removed using the Uchime algorithm and the no-chimeric sequences were clustered into OTUs at 97% identity using UPARSE software (20) (21). The RDP classifier was used to detect the taxonomic assignments of each OTU (22).

The merged sequences were submitted to the NCBI SRA database (accession number SRR10765290-SRR10765300).

### Data Analysis

The results for each parameter were presented as the mean ± standard error for each group. Principal coordinate analysis (PCoA) based on unweighted Unifrac distance was applied to evaluate the differences in different groups. Principal component analysis (PCA) was performed by R vegan package. Non-parametric ANOVA (PERMANOVA) was performed using the R vegan package(23) to analyze the significance of differences between groups; Welch’s T-test(STAMP) was performed to analyze significantly different phylum in gut microbiota between groups; Prism 6 was used for box plot production; T tests was conducted by SPSS16.0 to analyze the significance of differences in diversity indicators; the p-value less than 0.05 were significant difference, p-value less than 0.01 were significant extremely.

## Results

The skin of diseased *P. spinosa* had the white spot or large area of decay. After anatomy, there were obvious lesions in the liver and gallbladder. The liver blackened obviously. The gallbladder was enlarged or discolored. The symptoms of the diseased *P. spinosa* are shown in the picture (Fig. 1).

**Fig.1.**
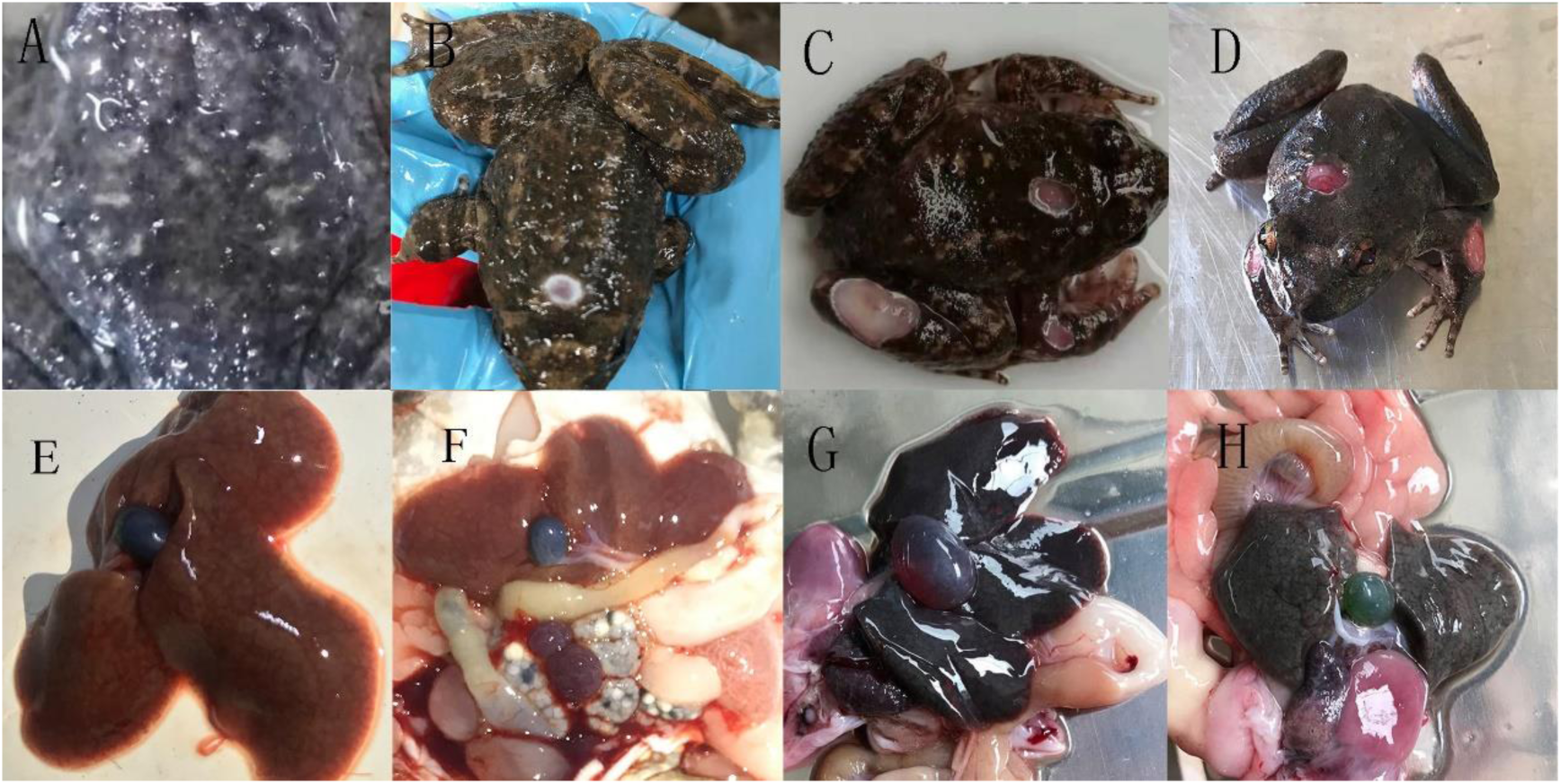
A and B are the frogs of the early stage of the rotten-skin disease; C and D are the later stages; E and F are the liver and gallbladder of healthy *P*.*spinosa*; G and H are the Lesions in the liver and gallbladder of diseased *P*.*spinosa*.

A total of 466,541 high-quality sequences were obtained from the 11 gut microbiota samples. To avoid the influence of sequencing depth, 25,520 sequences were randomly selected from each sample for further analysis, and 5,793 OTUs from 704 genera were identified. The results of gut microbiota diversity indicated that the diversity index, such as Chao1, Observed-species, PD-whole-tree, Shannon index, and Simpson index, in healthy *P. spinosa* were significantly lower than the diseased *P. spinosa*, but the Good-coverage had no difference (Fig.2). Removing the unclassified sequences (<0.001%),19 of the 56 phyla dominated the gut microbiota. *Bacteroides* and *Firmicutes* were the dominant microbiota in the gut of all samples (Fig. 3A), which is consistent with a previous study (1). Among them, the average relative abundance of *Bacteroides, Firmicutes, Proteobacteria, Tenericutes*, and *Euryarchaeota* was more than 1% in all samples (Fig. 3A). STAMP based on relative abundance of top 10 phyla in the gut microbiota showed that *Proteobacteria* was significantly higher in the healthy *P. spinosa*, while the relative abundance of *Euryarchaeota* and *Spirochaetes* were significantly lower (Fig. 3B).

**Fig.2.**
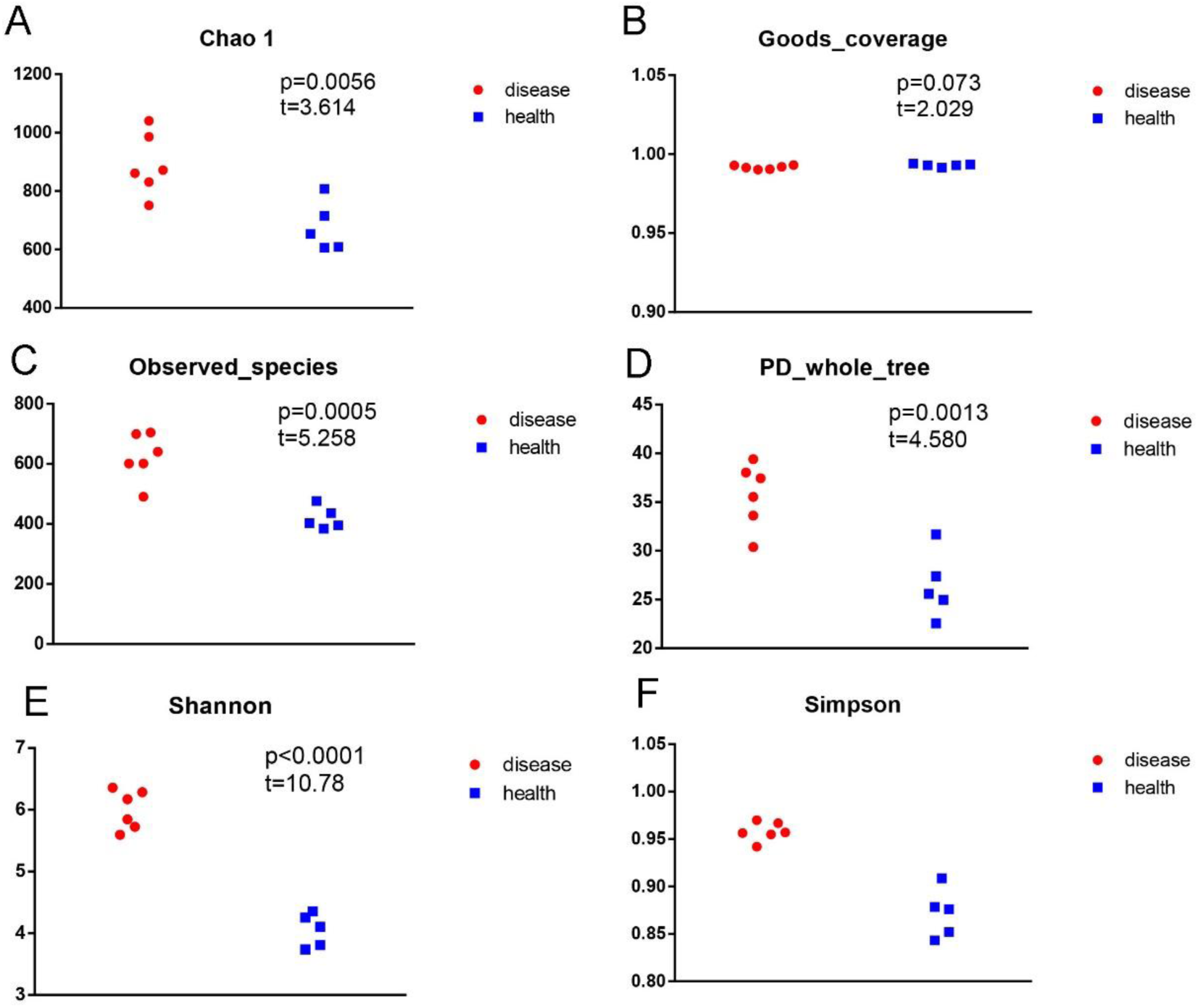
Diversity analysis of the gut microbiota *P. spinosa* between healthy and diseased groups: Chao1; (B) Goods-coverage; (C) observed-species; (D) PD-whole-tree; (E) Shannon index; (F) Simpson index. The gut microbiota of *P. spinosa* was collected from approximately 0.3g samples of the hindgut of each individual. Disease, the gut microbiota from six diseased *P. spinosa*. Health, the gut microbiota from five healthy *P. spinosa*. P values show the difference between the groups. P>0.05 represents the little difference between groups, p<0.05 indicates the significant difference between the groups, p<0.01 indicates the extremely different between the groups. Data are the mean± SE

**Fig.3.**
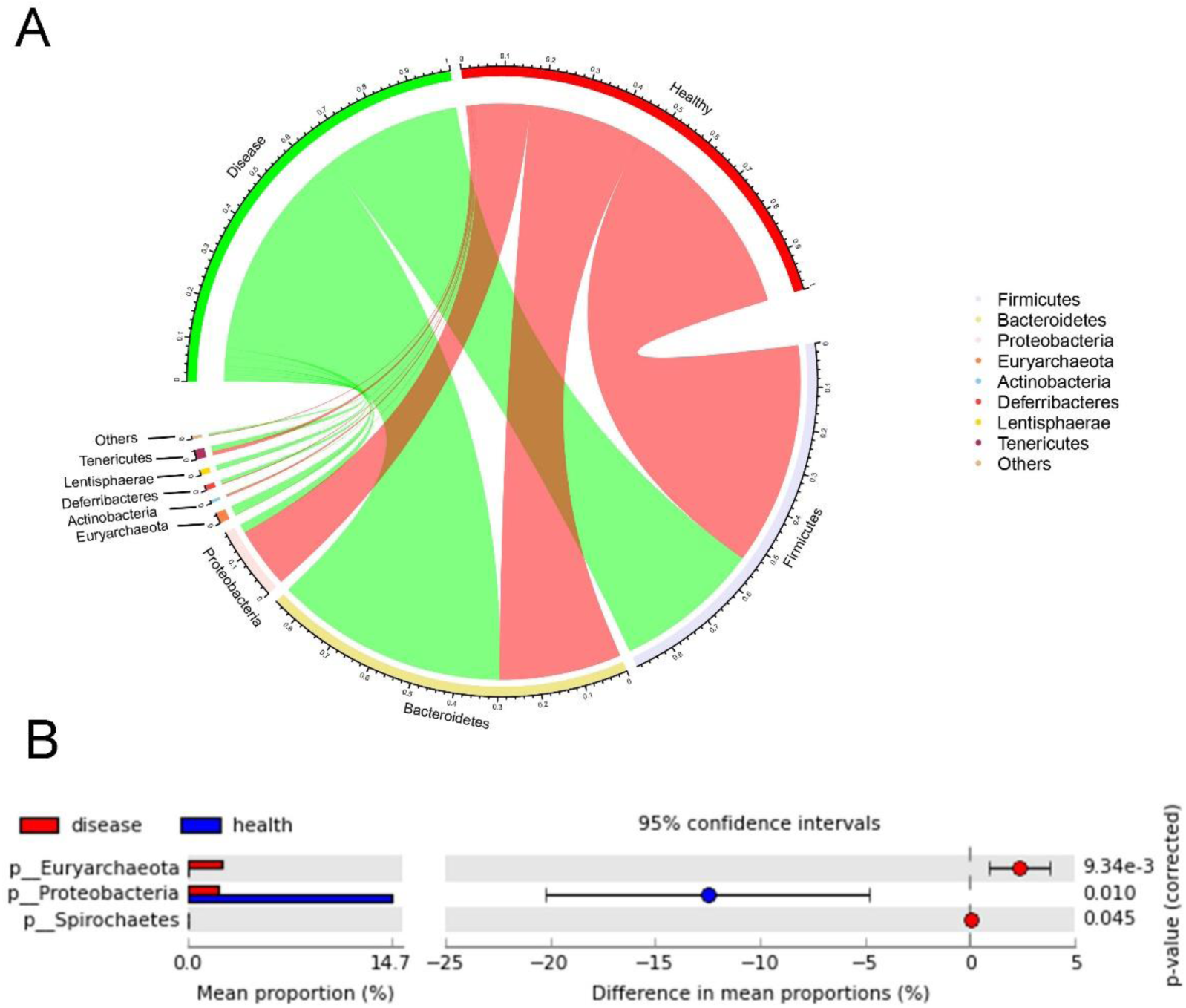
The inner circular(A) diagram shows the relative abundance of different phyla in *P*.*spinosa* gut samples. The gut microbiota of *P. spinosa* was collected from approximately 0.3g samples of the hindgut of each individual. D, the gut microbiota from six diseased *P. spinosa*. H, the gut microbiota from five healthy *P. spinosa*. the significant difference in phylum between health and disease groups(B). The STAMP based on the top 10 phyla of the gut microbiota compositions analyzed the significantly different(p<0.05) phylum between the groups.

PCA based on the relative abundance of all gut microbiota genera and PCoA based on the relative abundance of the all gut microbiota genera showed that there were significant differences in gut microbiota composition between diseased and healthy *P. spinosa* (PERMANOVA, *F*= 3.0464, *p* = 0.008) (Fig. 4A and 4B). Unweighted Pair-Group Method with Arithmetic means UPGMA analysis shown that microbiota composition were familiar between the groups(Fig. 4C).

**Fig.4.**
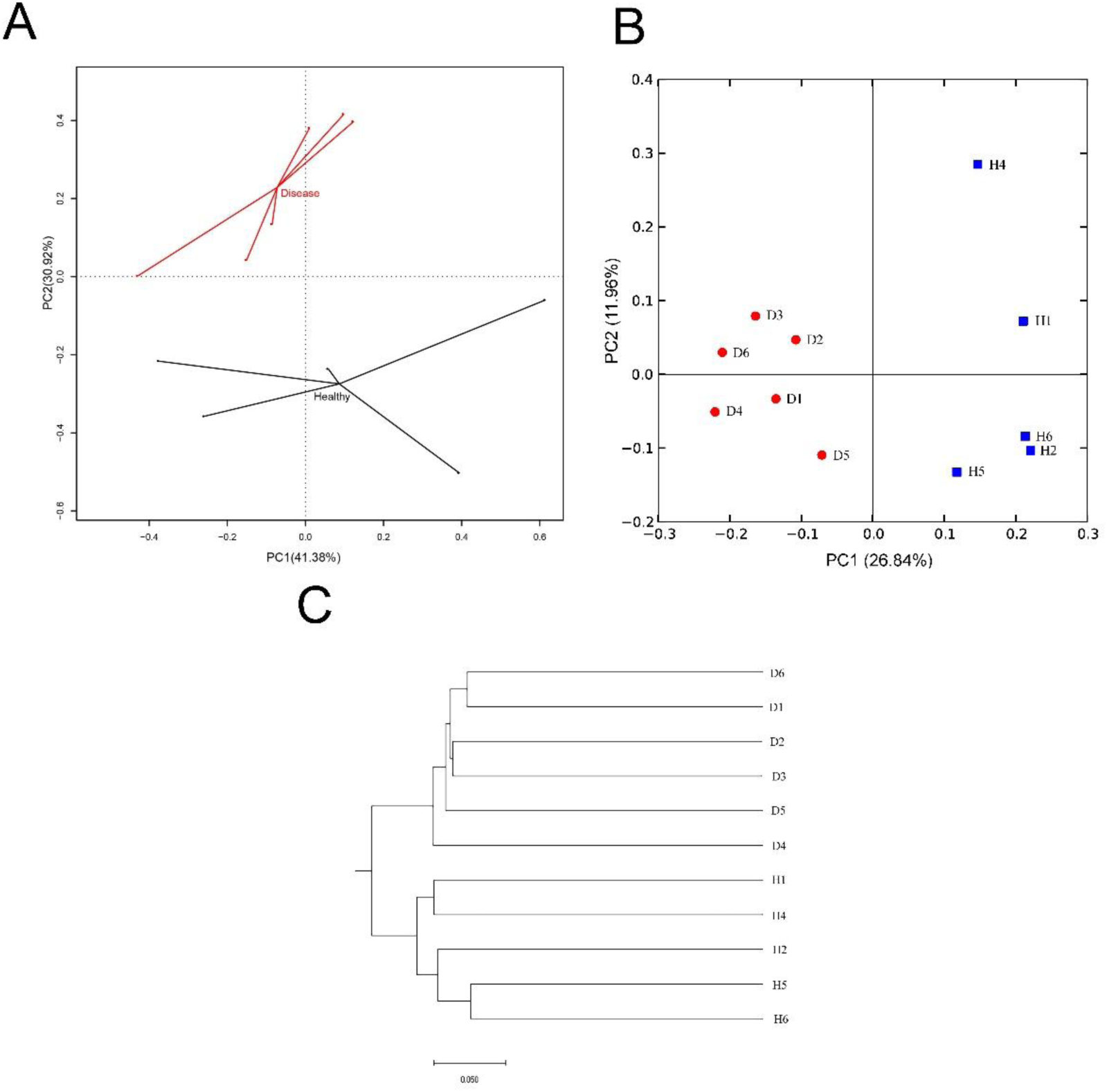
PCA profile(A). PCA was conducted based on the all genus microbial communities showing the differentiation of the *P*.*spinosa* gut microbiota communities between the health and disease group. PCoA profile(B)and UPGMA cluster graph (C) based on the unweight unifrac distance showing the differentiation of the *P*.*spinosa* gut microbiota communities between each individual. The PCoA was conducted based on the all genus microbial communities. The gut microbiota of *P. spinosa* was collected from approximately 0.3g samples of the hindgut of each individual. Disease, the gut microbiota from 6 diseased *P. spinosa*. Health, the gut microbiota from 5 healthy *P. spinosa*.

Lefse analyzed the difference of gut microbiota at genus level showed that relative abundance of *Serratia, Eubacteium, Citrobacter*, and *Morganella* were significantly higher in healthy *P. spinosa*, while the relative abundance of *Methanocorpusculum, Parabacteroides, AF12, PW3, Epulopiscium*, and *Oscillospira* were significantly lower (Fig. 5).

**Fig.5.**
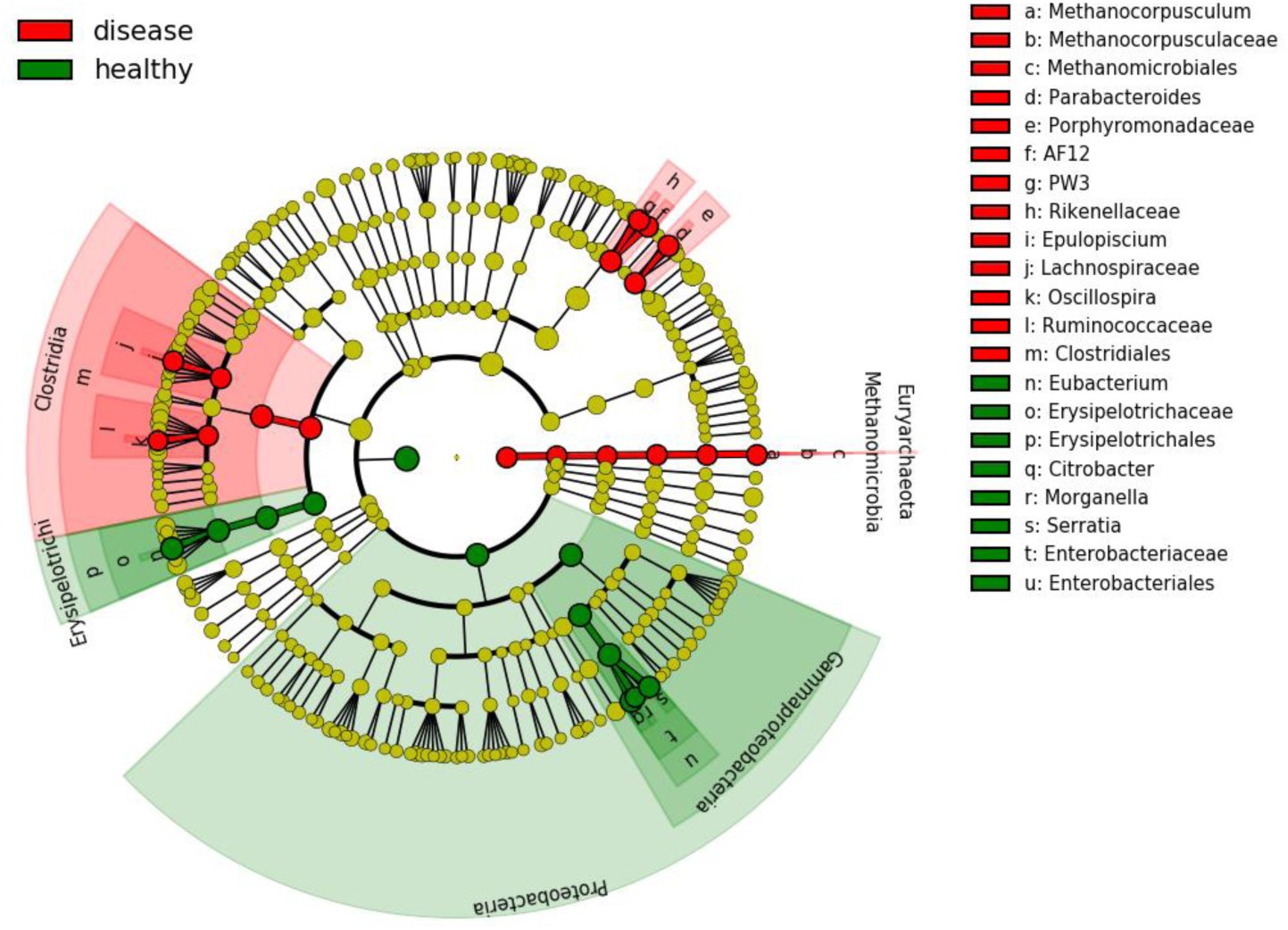
LEfSe profiles showing differences in healthy and diseased *P*.spinosa gut microbial communities. LEfSe analysis was conducted based on the top 40 genus compositions of the *P. spinosa* gut microbiota. The gut microbiota of *P. spinosa* was collected from approximately 0.3g samples of the hindgut of each individual. Disease, the gut microbiota from 6 diseased *P. spinosa*. Health, the gut microbiota from 5 healthy *P. spinosa*.

## Discussion

Recent researches have shown that gut microbiota has participated in various disease processes through the gut-brain axis (24) (25), the gut-lung axis (26) (27) the gut-vascular axis(28, 29), the gut-bone axis (30) (31),the gut-Hepatic axis (32) (33) and other axis (34). The concept of “core microbiota” indicated that the core microbiota in the gut of healthy hosts could maintain the stability of gut microbiota composition and function, and positively regulated the host through these axis to maintain host health (35). The gut microbiota function was destroyed because of the destruction of the core microbiota, and the host might become sick or aggravate the lesion (16). In this study, the gut microbiota of diseased and healthy *P. spinosa* was compared, and the results revealed significant differences in the gut microbiota composition of healthy and diseased *P. spinosa*. The composition of microbiota was destroyed because of pathogen invading.

Current researches on gut microbiota focused on gut microbiota diversity and gut microbiota composition. According to the diversity resistance hypothesis, the more diverse that the microbial community was and the more possible that the host resistanted to pathogen invasion (36). Studies in the largemouth bronze gudgeon (*Coreius guichenoti*) (37), crucian Carp (*Carassius auratus*) (14), and ayu (*Plecoglossus altivelis*) (38) showed that the microbiota diversity was significantly higher in healthy samples. However, this study found that gut microbiota diversity was significantly higher in diseased *P. spinosa*. The results were consistent with the results of grass carp (*Ctenopharyngodon idellus*) (39). And found that the amino acid metabolism, carbohydrate metabolism, and immune-related pathway genes of diseased grass carp were more abundant through microbiota gene prediction (39). The increased microbiota diversity in the gut of the diseased host may because the microbial homeostasis in the gut of the diseased host has not been broken immediately. To maintain the health of the host, the gut microbiota diversity was increased to protect against pathogen invasion. The results of this study and previous studies have shown that the use of gut microbiota diversity to assess host health is limited.

The relative abundance of *Methanocorpusculum, Parabacteroides, AF12, PW3, Epulopiscium*, and *Oscillospira* in the gut of rotten-skin *P*.*spinosa* were significantly higher than healthy *P. spinosa*. Although *Methanocorpusculum* is not a pathogen, it is abundant in diseased samples (40). It can effectively convert heavy metals or metalloids into more toxic derivatives than compounds, which was harmful to host health (41); *Parabacteroides goldsteinii* in *Parabacteroides* can cause bacteraemia; Previously studied in the gut microbiota of wild and cultured *P. spinosa* found that the cultured *P. spinosa* with more potential pathogens had more *AF12* in the gut (1); Studies on human gallstones indicated that the relative abundance of *Oscillospira* was positive correlation with the gallstones (42, 43); The relative abundance of *Epulopiscium* was significantly increased in the gut of rotten-skin diseased *P. spinosa*, and also significantly increased in the feces of children with eczema(44). In summary, the gut microbiota that was significantly increased in the gut of rotten-skin disease *P. spinosa* was mostly opportunistic pathogen; *Oscillospira* and *Epulopiscium* were significantly increased in the gut of the diseased host when lesions occurred in the skin and gallbladder. It was speculated that these two species may be indicator microbiota in the pathogenesis of skin and gallbladder.

The relative abundance of *Serratia, Eubacteium, Citrobacter*, and *Morganella* in the gut of healthy *P. spinosa* was significantly higher than diseased *P. spinosa*. Some species in *Serratia* produced Prodigiosin and β-lactam antibiotic carbapenem to inhibit the growth of pathogens in the host, thereby inhibiting the disease (45, 46); *Bifidobacteria* and *Eubacteium hallii* promoted acetate, butyrate, propionate, and formate to form, potentially contributing to gut SCFA formation with potential benefits for the host and for microbiota colonization of the infant gut (47). *E. hallii* was also capable of metabolizing glycerol to 3-hydroxypropionaldehyde with antibacterial properties (48); The relative abundance of *Citrobacter* and *Morganella* in the gut of healthy *P. spinosa* was significantly higher than diseased *P. spinosa*, however *Citrobacter rodentium* and *Morganella morganii* are common opportunistic pathogen (49) (50). In summary, *Serratia* and *Eubacterium* might be the main gut microbiota in the healthy *P. spinosa* that maintained the health of *P. spinosa*; *Citrobacter* and *Morganella* in the gut of healthy *P. spinosa* were significantly increased without causing disease. It might be that non-pathogenic strains of *Citrobacter* and *Morganella* appeared in the gut of healthy frogs, or there were pathogenic strains in these two species, but due to the inhibition of beneficial microbiota in the gut of healthy hosts, the host still maintained a healthy state of gut microbiota homeostasis.

## Conclusion

Rotten-skin disease significantly changed *P. spinosa* gut microbiota; the relative abundance of *Methanocorpusculum, Parabacteroides, AF12, PW3, Epulopiscium*, and *Oscillospira* were significantly higher in the diseased *P. spinosa*, while the relative abundance of *Serratia, Eubacteium, Citrobacter*, and *Morganella* were significantly lower; The relative abundance of *Epulopiscium* and *Oscillospira* might be related to the healthy condition of the host skin and gallbladder; The relative abundance of *Serratia* and *Eubacteium* might be important for maintaining the gut microbiota ecosystem.

